# Minimizing command timing volatility is a key factor in skilled actions

**DOI:** 10.1101/2024.06.11.598574

**Authors:** Atsushi Takagi, Sho Ito, Hiroaki Gomi

**Affiliations:** NTT Communication Science Laboratories, 3-1 Morinosato Wakamiya, Atsugi, Kanagawa, 243-0198, Japan

## Abstract

Variability between movements prevents the best athletes from making a perfect shot every time. While fluctuations in the amplitude of neural sensory inputs and motor outputs are thought to be primarily responsible, they only account for a fraction of the observed variability. Here, we propose that a significant portion of the variability is due to imprecisely timed motor commands. This command timing volatility theory best explained the three peaks observed in the force variability’s time-series in discrete reaching movements and during periodic force control. Furthermore, we show how the timing volatility in the non-dominant arm’s muscles is larger than in the dominant arm, then develop a variability index that estimates the arm’s timing volatility via its variability during circle tracing. The difference in the variability index between the left and right hands accurately predicts the Edinburgh Quotient, suggesting a relationship between handedness and the command timing volatility of the left- and right-hands. Lastly, we constructed a simulation of reaching movements made by an arm controlled by muscles whose command timing was made incrementally more volatile. As timing volatility increased, aiming became less precise and movements jerkier. Such impairments during reaching are reported in patients with different neuronal diseases that damage any brain regions critical to motor timing, suggesting that essential aspects of these symptoms may be caused by excessive timing volatility. Our theory provides a unifying computational perspective of movement variability in healthy and diseased individuals that is essential to understanding the control of movements.

## Introduction

Athletes and musicians marvel us with expert skills that are speedy yet precise. We have ideas concerning how the brain learns to speed-up movements using chunking (Rhodes et al., 2004), efficient neural representations (Krakauer et al., 2019) or preparation (Zimnik and Churchland, 2021), but the process of reducing movement variability with practice is not well understood and is understudied (Shmuelof et al., 2012). One contributor to movement variability is peripheral signal-dependent noise (SDN) in the muscles, whose amplitude increases with its activation size (Harris and Wolpert, 1998). Sensory noise during the movement planning stage can add to movement variability too in the form of spatial noise (SN), where misestimation of the distance between the target and hand in hand-centered coordinates can occur (Gordon et al., 1994). Another contributor to movement variability is temporal noise (TN) during movement planning, which causes fluctuations in the planned movement duration that distort both the amplitude and the duration of the planned muscle activity between movements (Meyer et al., 1982; van Beers et al., 2004).

Recent findings, however, call into question whether these sources fully account for movement variability. The proportion of SDN relative to the amplitude of the exerted force is empirically measured to be approximately 2% (Adam et al., 1998; Jones et al., 2002), but at least 20% is needed to produce human-like movement variability in simulations of reaching movements (Berret and Jean, 2020; van Beers et al., 2004). While SN does influence the variability of movements, its effect is too minor to explain movement variability when both the target and the hand are clearly visible (Beers et al., 1999; Hansen and Skavenski, 1977). And while TN does increase movement variability, its contribution is shown to be smaller than SDN’s in simulated reaching movements (van Beers et al., 2004). Therefore, signal-dependent noise, spatial noise, and temporal noise together cannot account for a significant portion of the variability between movements. Here, we propose an alternative, central source of movement variability that is determined by fluctuations in the timing of the commands that activate each muscle.

First, we compare the predictions driven by the existing theories and the new CTV theory. Then, three experiments will be introduced to examine these predictions. The first experiment tested discrete elbow reaching movements to compare the measured force variability against the predictions of the four theories. Next, we examined the force variability during periodic force control to confirm whether the force variability during periodic actions is best described by CTV theory. Then, we developed a variability index where the left and right arm’s timing volatility is estimated via its variance during circle tracing, and in a third experiment show how the difference in the variability index between the hands is related to handedness. Finally, we constructed a simulation of reaching movements using an arm controlled by muscles whose timing was made more volatile. This simulation shows how the motor symptoms caused by increased command timing volatility qualitatively resemble the problems in aiming and excessive movement jerkiness that patients with Huntington’s disease (Smith et al., 2000), Parkinson’s disease (Sheridan et al., 1987), and cerebellar disorders exhibit (Sanguineti et al., 2003), thereby supporting our hypothesis that precise command timing of the muscles is a key aspect of coordinating movements.

### Predictions of existing and new theories

To compare the difference between the newly proposed CTV theory relative to SDN, SN, and TN, let us consider an example of moving and stopping a single joint in one smooth movement using two muscles (see Methods for details). In this example, the agonist minus the antagonist activity is assumed to be equal to the force the joint produces, i.e., each muscle can “cancel out” the other’s force because they pull in opposite directions. When moving and stopping the joint, the agonist provides the accelerative force and the antagonist produces the braking force (**Figure 1A**). Here, we introduce the command timing volatility theory, which proposes that noise corrupts the activation timing of each muscle. If a muscle is activated at the unintended time, the force time-series deviates from the planned or ideal one, thereby causing force variability. Since position is the double integral of the force time-series, any variation in the force will lead to position variability too. The SDN, SN, TN, and CTV theories generate different predictions concerning how the force variability of 200 simulated movements unfolds with time as shown in **Figure 1A**.

**Figure 1.**
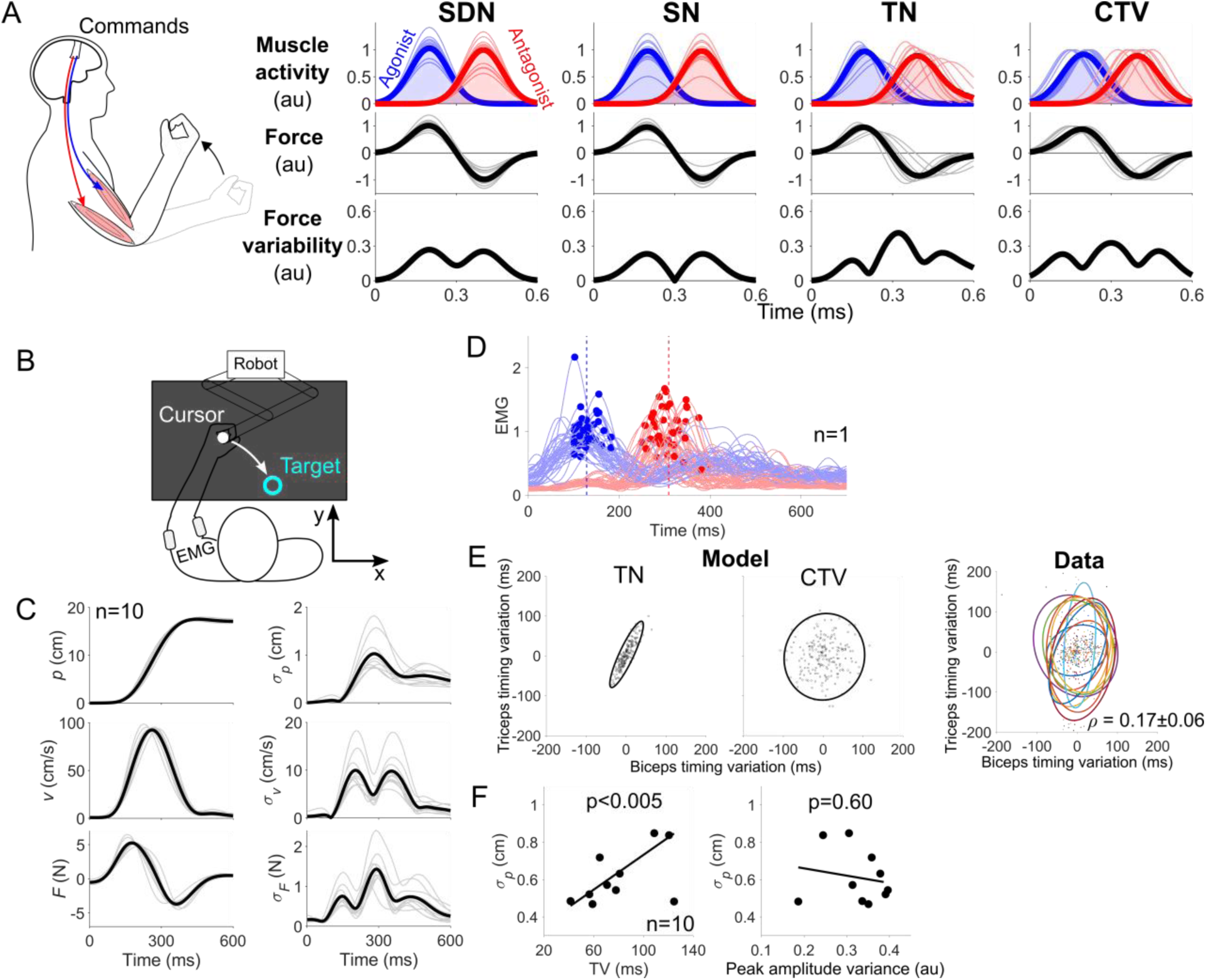
Movement variability primarily comes from command timing volatility. (A) SDN and SN theories predict two peaks in the force-variability time-series, while TN and CTV predict three peaks centered at the change in force direction. (B) Schematic of elbow reaching experiment. (C) Time course of the elbow’s position *p*, velocity *v* and force *F* exerted against the handle and their respective variability (σ*_p_*, σ*_v_* and σ*_F_*). The force variability exhibited three peaks. (D) Rectified and smoothed biceps brachii and triceps lateral head activity aligned at movement onset from a representative participant. Variability in the amplitude and timing of the activity’s peak was observed. (E) TN theory predicts high correlation between the timing variation in the two muscles or a narrow covariance ellipse along the diagonal (left). CTV theory predicts low correlation or a round covariance ellipse (middle). Actual data showed a low correlation in the muscles’ timing variation (right, each color denotes a different participant). (F) TV was linearly related to the elbow’s position variability (left panel), but the variance in peak EMG amplitude was not (right panel).

The SDN theory predicts variations in the amplitude of the muscle activity. This was simulated by multiplying each muscle’s activity time-series by a scaling factor corrupted by noise that was separate for each muscle and which was varied between movements. The SDN theory predicts that the force should vary the most where the muscle activity is greatest, resulting in two large peaks in the force variability time-series **(Figure 1A, left panels)**. The SN theory considers noise in planning the length of the movement. This was simulated by multiplying both muscles’ activity by the same scaling factor that was corrupted by noise. The SN theory predicts two peaks in the force variability time-series **(Figure 1A, panels second from left)**. The TN theory considers noise in the total duration of the movement. This is simulated by scaling the muscle activity time-series by a time-scaling factor that was corrupted by noise, and then by multiplying the muscle activity by the inverse of the time-scaling factor. This preserves the area under the muscle activity curve, which ensures that the joint moves to the same location in space but at different times. This causes common shifts in the timing of the peak agonist and antagonist muscle activity, so their timing should be strongly correlated. Interestingly, the TN theory predicts three peaks in the force variability time-series **(Figure 1A, panels second from right)**. The CTV theory proposes that there is fluctuation in the activation timing of the agonist and antagonist muscles that is independent of each other. This was simulated by adding separate noise to the timing of the agonist and antagonist muscles between movements. The CTV theory predicts three peaks in the force variability, but the timing of the peak agonist and antagonist activity are weakly correlated **(Figure 1A, right panels)**. Does human force variability unfold according to the SDN, SN, TN, or CTV theory?

## Results

### Position variability is determined by timing volatility

To test the predictions of each theory, we examined the force variability when healthy participants (n=10) made reaching movements towards a target with their left elbow (**Figure 1B**). The elbow’s trajectory followed the classically observed bell-shaped velocity time-series (**Figure 1C**, left middle panel). We then analyzed how the variability in the elbow’s position, velocity, and the force exerted against the handle (σ*_p_*, σ*_v_* and σ*_F_*) unfolded along the movement. The force variability exhibited three peaks (**Figure 1C**, bottom-right), which is inconsistent with the predictions of SDN and SN theories but is consistent with the TN and CTV theories shown in **Figure 1A**. Although the TN and CTV theories are not distinguishable in terms of the force variability time-series, TN predicts a strong correlation between the timing of peak agonist and antagonist muscle activity. The CTV theory, on the other hand, expects low correlation in the peak timing of the two muscles since the noise affecting each muscle’s command timing is independent for each muscle.

To differentiate between the TN and CTV theories, we examined temporal relationship between the activities of the agonist and antagonist muscles. We rectified and smoothed the electromyogram (EMG) of the biceps brachii (agonist) and the triceps lateral head muscles (antagonist), then calculated the timing of the peak agonist and antagonist activities in every trial, which varied relative to movement onset (**Figure 1D**, representative participant). We subtracted the mean from each muscle’s peak timing to examine the correlation in the timing variation between the agonist and antagonist muscles. According to the TN theory, when the variation in the peak timing of the agonist is plotted against the variation in the antagonist’s peak timing, a narrow covariance ellipse oriented along the diagonal should be visible as the timing variations should be correlated since both muscles are affected by the same timing noise factor (**Figure 1E**, left, 200 simulated movements). On the other hand, the CTV theory predicts a round covariance ellipse because the command timing to each muscle is corrupted by independent noise (**Figure 1E**, middle). The timing variations from the biceps and triceps muscles revealed a round covariance ellipse (**Figure 1E**, right) whose correlation was low at *ρ* =0.17±0.06 (mean±SEM), which is more consistent with the CTV theory. Thus, the three peaks in the force variability time-series likely originated from the volatile timing of the biceps and triceps muscles.

We then examined whether the timing volatility of the elbow muscles could predict the elbow’s position variability. The standard deviation of the peak timing relative to movement onset was calculated separately for each muscle as a measure of its timing volatility (TV). We then calculated the elbow’s mean position in the time window starting from 150 ms to 600 ms after movement onset for each trial, then calculated the standard deviation of this position variability across all 50 trials, which is defined to be σ*_p_*. This position variability was plotted as a function of the mean TV of the biceps and triceps muscles (**Figure 1F**, left). Robust regression of these two variables revealed a significant linear relationship between them (slope=0.005±0.001 mean and standard error, p=0.005), suggesting that movements controlled by muscles with larger TV lead to larger position variability.

While the peak timing of the biceps and triceps muscles varied between movements, we also observed significant variance in the peak amplitude of the muscle activity during elbow reaching (**Figure 1D**). To see if the variance in muscle activity amplitude could also explain the elbow’s position variability, we calculated the variance in the biceps and triceps’ peak EMG amplitude, averaged these values then plotted it as function of the elbow’s position variability (**Figure 1F**, right). A linear regression analysis revealed no significant relationship between the variance in the EMG’s peak amplitude and the elbow’s position variability (slope=-0.46±0.84, p=0.60). Thus, the elbow’s position variability cannot be attributed to variations in the muscle activity’s amplitude. Instead, it appears to be linearly related to the timing volatility of the muscles controlling the movement.

### Timing volatility of the muscles in the dominant and non-dominant arms

In the first experiment, we showed how the timing volatility of the elbow muscles is linearly related to the elbow’s position variability during discrete reaching movements. The aim of the second experiment was to extend this finding to periodic actions in different joints in the left and right arms. In the second experiment, right-handed participants (n=10) periodically activated the agonist and antagonist muscles of either the wrist, the elbow, or the shoulder against a fixed handle that was equipped with a force sensor. When testing one joint, e.g., the wrist, the other two joints were fixed by a brace to prevent their motion from affecting the force readings (**Figure 2A**). The muscle activity from two monoarticular muscles spanning each joint were filtered and normalized to have a mean peak activity equal to one.

**Figure 2.**
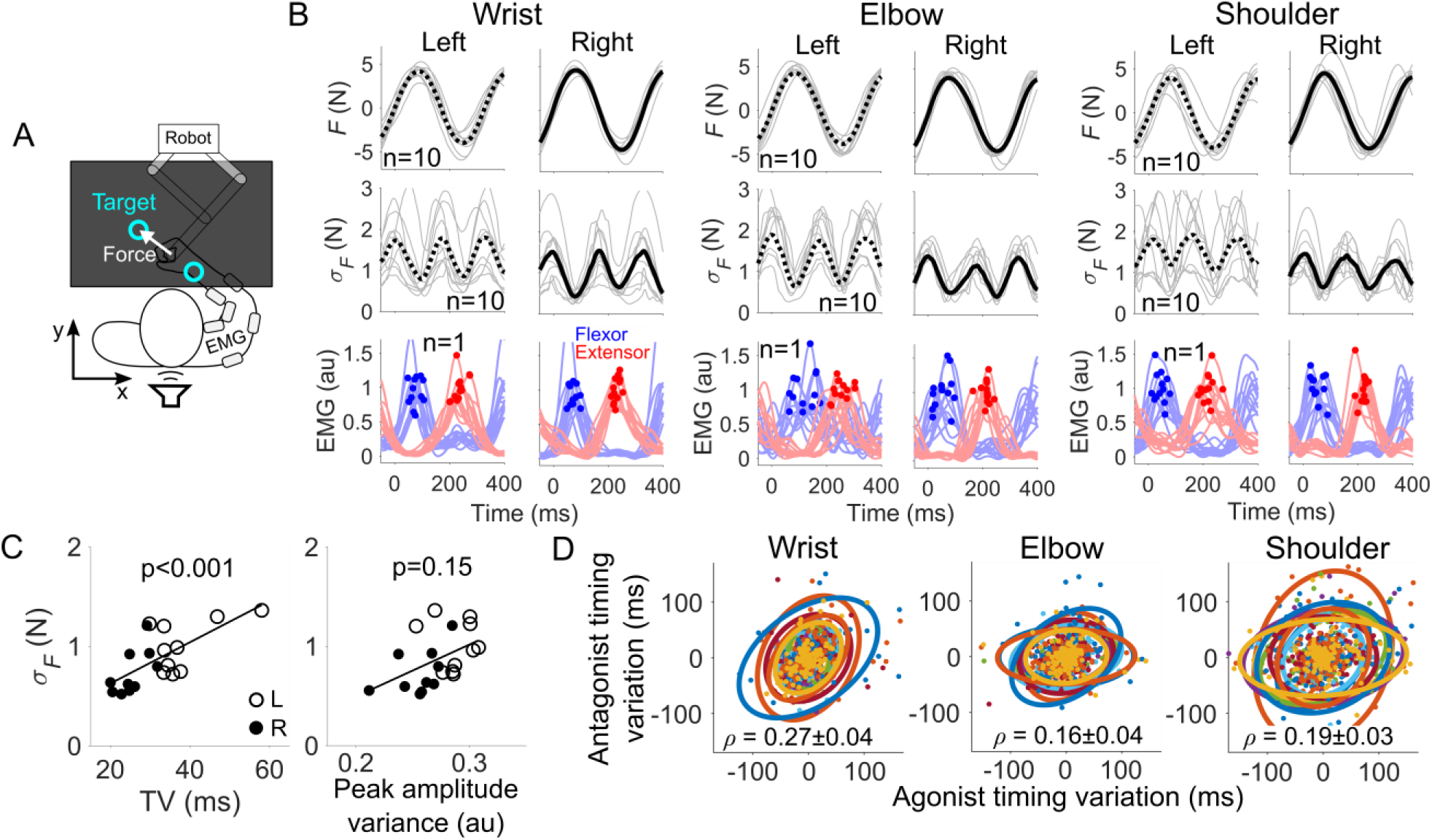
Timing volatility is different between the left and right arm’s muscles. (A) Schematic of the experiment to measure the TV of the muscles isometrically. (B) Time-series of the mean force and force variability from the wrist, elbow, and shoulder from all participants (n=10). Time-series of the muscle activity is shown from the same exemplar participant (n=1). Force variability had three peaks and the timing of peak muscle activity, marked by colored dots (flexor in blue, extensor in red), was more variable in the left arm. (C) Mean TV, averaged from all six muscles, was linearly related to the mean force variability, but variance in peak EMG amplitude was not (open circles denote the left arm, filled are the right arm). (D) Variation in antagonist’s peak timing versus agonist peak timing variation from the wrist, elbow, and shoulder joints. The correlation between the two timing variations was low in all three joints, which is consistent with CTV theory.

The force time-series was separated into a push-pull cycle such that the agonist’s activation preceded the antagonist’s (**Figure 2B**). The force variability time-series of the wrist, elbow, and shoulder all had three peaks, which is consistent with the TN and CTV theories (**Figure 2B**, second row of panels). The activity of the biceps and triceps muscles from a representative participant show how the muscles’ TV appeared to be larger in the left elbow than in the right one (**Figure 2B**, bottom panels). To quantify this difference, the TV of the agonist and antagonist muscles were calculated by taking the timing difference between the preceding and proceeding peaks, and then by calculating the standard deviation in this timing difference across all trials. The force variability (t(9)=4.39, p=0.002), the TV (t(9)=4.26, p=0.002), and the variations in peak EMG amplitude (t(9)=3.49, p=0.007) from all participants across all three joints were all larger in the left arm (post-hoc tests controlled for multiple comparisons using the Holm-Bonferroni method).

Why is the force variability larger in the non-dominant left arm? The larger force variability in the non-dominant arm cannot be explained by the SN theory since the force variability time-series had three peaks and not two. While this also discounts the SDN theory, the variance in peak EMG amplitude was larger in the left hand, so the SDN theory is still plausible. And the TN and CTV theories could explain the force variability because they both predict three peaks in the force variability time-series. Which theory best explains the force variability during periodic actions?

We conducted a linear mixed-effects analysis with the force variability as the dependent variable and the hand (left, right) and the joint (wrist, elbow, shoulder) as categorical predictor variables, and the TV and the variance in peak EMG amplitude as continuous predictor variables. The joint predictor did not improve the fit of the linear model (*χ*(2)=1.2, p=0.55) (see **Figure S1** in Supplementary Materials for fits separated for each joint), so the values for the TV and peak EMG amplitude variance from the muscles in the same arm were averaged in **Figure 2C**. Further analysis using the linear mixed-effects model revealed a significant linear relationship between the TV and force variability (**Figure 2C**, left, *χ*(1)=11.5, p<0.001), but variance in peak EMG amplitude was not significantly related to the force variability (**Figure 2C**, right, *χ*(1)=2.1, p=0.15). While variations in peak EMG amplitude were overall larger in the left hand, larger EMG amplitude variance did not imply larger force variability. Therefore, the SDN theory cannot explain why the non-dominant arm’s force variability was greater.

This leaves the TN and CTV theories as plausible sources for the force variability. To distinguish between them, we analyzed the correlation in the timing variation between the agonist and the antagonist muscles from the wrist, elbow, and shoulder joints separately (**Figure 2D**). The timing variation for each muscle was calculated by obtaining the mean timing interval between the peaks in muscle activity in one trial, then taking the difference between this mean timing interval and the actual timing interval between neighboring peaks in muscle activity. This was carried out separately for each muscle in every trial to get the plot in **Figure 2D**. The TN theory expects a strong correlation in the timing variation of the agonist and antagonist muscles. When plotted against each other, the TN theory predicts a narrow covariance ellipse oriented towards the diagonal, as examined earlier in the first experiment (**Figure 1E**, left), but the actual covariance ellipses were round as CTV theory expects (**Figure 1E**, right). The correlation values were *ρ* =0.27±0.04, *ρ* =0.16±0.04, and *ρ* =0.19±0.03 for the wrist, elbow, and shoulder joints, respectively. This suggests that the CTV theory best explains why the non-dominant wrist, elbow, and shoulder produce a force that is more variable than those exerted by the dominant arm.

Based on the results of the second experiment, we hypothesized that the timing volatility of the left and right arm’s muscles could be related to the handedness of the participants. To confirm our hypothesis, we developed a variability index that relates the arm’s average timing volatility, taken from the wrist, elbow, and shoulder muscles, to the arm’s periodic movement variability. This variability index allowed us to estimate the gross timing volatility of the muscles in the left and right arms sans EMG. The variability index was measured by recording the hand’s acceleration as it traced a circle of diameter 10 cm unimanually for 15 seconds at a speed of 2.5 Hz (**Figure 3A**). First, we collected data from the people who had participated in the second isometric force control experiment where EMG timing volatility data was also available. The acceleration traces from the left and right hands were scaled for comparisons, which showed that the left-hand’s trace was more variable **(Figure 3B)**.

**Figure 3.**
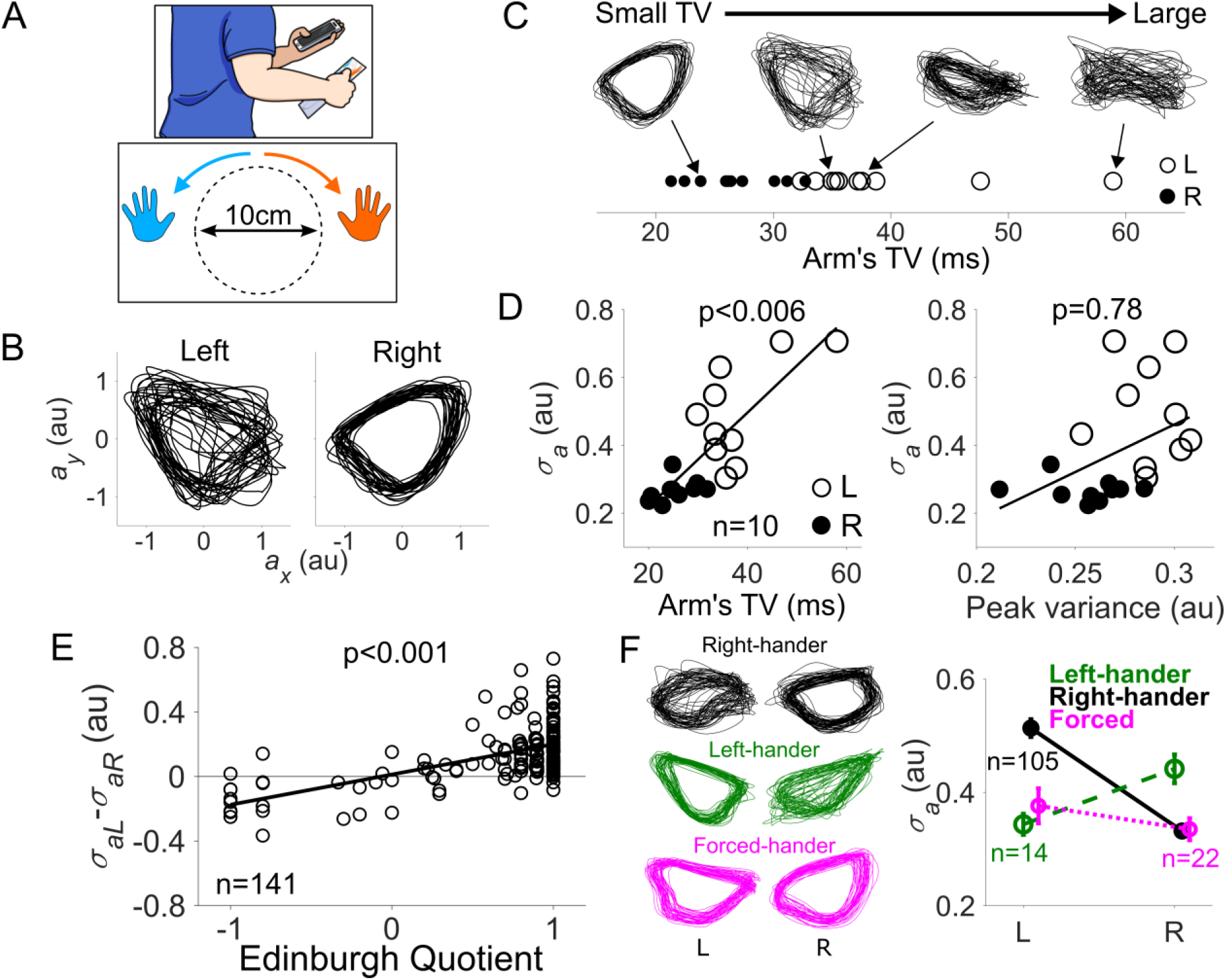
The dominant hand is empowered by small TV. (A) Schematic of the experiment to measure periodic movement variability when drawing circles at 2.5 Hz with a smartphone. (B) Acceleration trajectories from the left and right hands of a representative participant (only *x* and *y* axes shown for visibility) showed greater movement variability in the non-dominant left hand. (C) Arm’s TV or the average TV of the six muscles measured from the isometric force control experiment (open circles are from the left arm, filled from the right arm), and the corresponding arm’s periodic acceleration trajectory from four representative participants. When the arm’s muscles had large TV as estimated from the EMG, the periodic acceleration variability was also greater. (D) Arm’s TV was a reliable predictor of the hand’s acceleration’s variability σ*_a_*, but variations in peak EMG amplitude was not. (E) Left acceleration variability σ*_aL_* minus right acceleration variability σ*_aR_* was linearly related to the Edinburgh Quotient. (F) Forced-hander’s right-hand had smaller acceleration variability relative to left-handers, demonstrating how prolonged motor training may reduce the arm’s TV.

To see if the acceleration variability is related to the timing volatility of the arm’s muscles, we calculated the TV of the arm, which is defined to be the mean timing volatility from the six muscles of the arm (wrist, elbow, and shoulder flexor and extensors) from the second experiment. The TV of the arm is charted on the horizontal axis alongside the acceleration trace from four representative participants **(Figure 3C)**. Upon visual inspection, the arm whose muscles had larger TV in the second experiment exhibited acceleration traces that were more variable in the third experiment. To quantify this relationship, we estimated the variability of the acceleration trace by separating each cycle of the circular trace, then estimated the similarity of the two neighboring circular traces by calculating the Hausdorff distance between them, a commonly used shape similarity metric (Huttenlocher et al., 1993). For a 15 second movement at 2.5 Hz, this yielded roughly 37 circular traces, and then 36 Hausdorff distances were calculated and averaged to yield the acceleration variability σ*_a_*. We then plotted σ*_a_* as a function of the arm’s TV (**Figure 3D**, left), and used a linear mixed-effects analysis with σ*_a_* as the dependent variable and the categorial hand variable (left, right), the arm’s TV as predictors, which revealed a significant linear relationship between the acceleration variability and the arm’s TV (*χ*(1)=7.6, p=0.006). Thus, the TV of the arm’s muscles is positively related to the arm’s variability when tracing circles. We also checked whether σ*_a_* could be explained by the average variance in the muscle’s peak EMG amplitude averaged across all six muscles (**Figure 3D**, right), but the linear mixed-effects analysis revealed that this relationship was not significant (*χ*(1)=0.08, p=0.78). Thus, the variability during circular tracing in the third experiment could be explained by the TV of the six muscles in the arm measured in the second experiment, but the variance in the peak EMG amplitude of these same muscles was not related to the same arm’s acceleration variability.

Using this acceleration variability index σ*_a_*, we asked 141 newly recruited participants to trace circles with a smartphone using their left and right hands (**Figure 3A**). The handedness of these participants was assessed using the Edinburgh Handedness Inventory (Oldfield, 1971), which revealed that 14 were left-handed (Edinburgh Quotient less than zero) and 105 were right-handed (Edinburgh Quotient greater or equal to zero). The remaining 22 were left-handers who were coerced into using their right hand, a practice known as forced correction, who are henceforth referred to as forced-handers.

If the TV of the arm’s muscles, which is related to σ*_a_*, is related to the laterality of the hands as we hypothesized, then a relationship should exist between the difference in the left and right hand’s acceleration variability σ*_aL_* − σ*_aR_* and the Edinburgh score. We charted σ*_aL_* − σ*_aR_* as a function of the Edinburgh Quotient (**Figure 3E**), and a regression analysis revealed that these two variables were significantly related (slope=0.16±0.02 mean and standard error, p<0.001).

We also looked at the acceleration variability of the left-hand σ*_aL_* and the right-hand σ*_aR_* separately for the left-handers, right-handers, and forced-handers. Forced-handers’ acceleration variability appeared to be small in both the left- and right-hands (**Figure 3E**). To see if the acceleration variability was different between the left- and right-hands of left-handers, right-handers, and forced-handers, we conducted a two-way repeated measures ANOVA with the acceleration variability as the dependent variable and the categorial hand (left, right) and categorial group (left-hander, right-hander, forced-hander) variables as the main predictors. σ*_aL_* and σ*_aR_* were significantly different within left-handers (p<0.001), within right-handers (p<0.001), and within forced-handers (p<0.001). However, the σ*_aL_* of forced- and left-handers was not significantly different (p=0.83), and neither was σ*_aR_* of forced- and right-handers (p=0.96), meaning that forced-handers could trace circles with the left and right hands equally well as the dominant hand of left-handers and right-handers, respectively, making them ‘doubly dominant’. The forced-hander’s right-hand variability σ*_aR_* should have been as large as the σ*_aR_* of left-handers since they were born as left-handers, but it was significantly smaller (p<0.001). This suggests that prolonged motor training of the right-hand reduced the right-hand’s acceleration variability, possibly by a reduction in the TV of the right arm’s muscles.

### Simulated increase in timing volatility breaks down motor coordination

The second experiment showed how the timing volatility in the muscles caused larger force variability during a periodic timing task. What happens to movement coordination as timing volatility increases to excessive levels? When patients with Huntington’s disease (HD), Parkinson’s disease (PD), and cerebellar ataxia (CA) do a periodic tapping task, they exhibit larger variability in the timing between taps relative to age-matched controls (Michell et al., 2008; O’Boyle et al., 1996; Schlerf et al., 2007). This raises the possibility that HD, PD, and CA damage different parts of the brain that are critical to motor timing, which may lead to larger TV. Can some of the motor symptoms shared by patients with HD, PD and CA be predicted by the CTV theory?

To explore this possibility, we conducted simulations of reaching movements using a computer simulation of a brain controlling a two-joint arm using six muscles (see Methods). Using this model, we simulated movements in eight directions with a healthy TV of 80 ms, which was the average measured in the first experiment, and then observed the consequences of increasing TV by 100%, 200% and 300%, corresponding to TVs of 160, 240, and 320 ms. For reference, HD patients have 20-400% greater timing variability relative to controls (Michell et al., 2008), and PD and CA patients exhibit approximately 60-70% greater timing variability (O’Boyle et al., 1996; Schlerf et al., 2007). 100 movements were simulated per reaching direction for each level of TV.

When the TV was 80 ms, the simulated hand’s position moved in a straight-line and stopped inside the target, but its trajectory became less straight and jerkier as TV increased (**Figure 4A**, top panels, only two sample trajectories shown). With TV of 320 ms, the simulated hand occasionally did not reach the target at all. To quantify the jerkiness of the trajectory, we calculated the mean-squared jerk (the derivative of the acceleration), as a function of time separately for the four (80, 160, 20, 320 ms) TV values (**Figure 4A**, bottom panel). The mean-squared jerk was the same for all values of TV in the early phase of the movement, but after 500 ms the jerk was greater when TV was larger. This difference in jerk resembles the reaching movements of patients with HD who exhibit larger jerk towards the end of the motion relative to age-matched controls (Smith et al., 2000). This suggests that excessive TV may impair the control process needed to reach straight and stop at a target location successfully.

**Figure 4.**
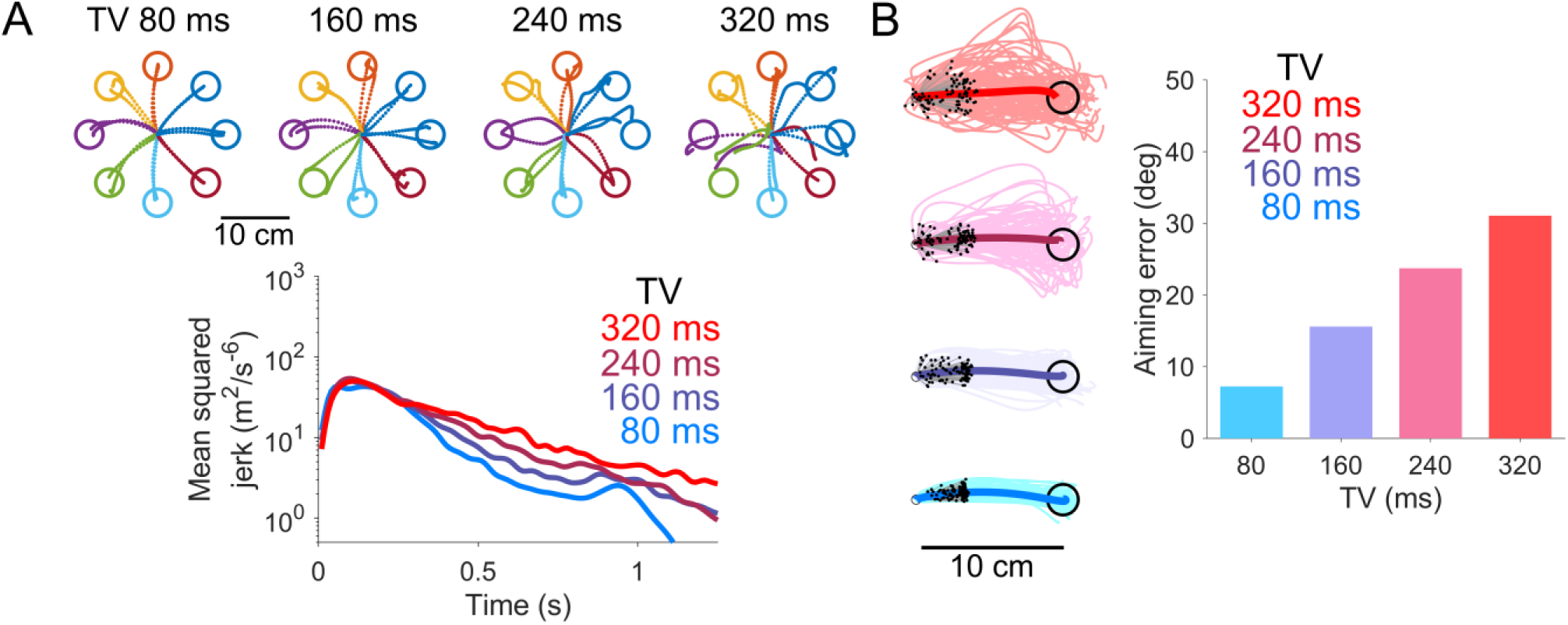
Extremely volatile command timing causes jerkiness and impaired aiming during simulated reaching movements. (A) Simulation of planar reaching to eight equidistant targets with TV of 80 (healthy), 160, 240, and 320 ms (two sample trajectories shown per target). Trajectories were less straight and jerkier at movement termination with increasing TV (bottom). (B) 100 simulated movements to the same target with TV of 80, 160, 240 and 320 ms. Thick trace is the mean trajectory. Aiming error was calculated 300 ms after movement onset before peak movement velocity (black dots). Aiming error increased significantly with larger TV (right). The details of the simulation dynamics are described in the Methods subsection titled ‘*Simulation of the brain controlling a two joint arm’*.

While increasing the muscles’ TV did not increase the jerk at the beginning of the movement, the simulated hand appeared to have problems aiming as TV was increased. To visualize the aiming error, we plotted all 100 trajectories when reaching towards the same direction with TV of 80, 160, 240, and 320 ms, and labelled the simulated hand’s position 300 milliseconds after movement onset (**Figure 4B**, black spots in left panels). As the simulated muscles’ TV increased, the hand’s position dispersed along directions orthogonal to the target. We quantified this dispersal by calculating the aiming error, defined as the absolute difference in the target direction and the direction of the hand’s position 300 ms after the beginning of the movement. The standard deviation of the aiming error increased monotonically with TV (**Figure 4B**, right panel). These simulation results suggest that increasing TV not only causes movements to be jerkier, but it impairs aiming when reaching towards a target as well.

## Discussion

A detailed examination of the muscle activity’s peak timing and its relationship with the force variability during discrete movements and periodic actions supports our hypothesis that central command timing volatility is a significant factor behind movement variability. Variability between movements has been attributed to wrongly-sized commands that stem from fluctuations in the amplitude and duration of the movement plan and to noise in movement execution. Movement variability can also be shaped by the interaction between the dynamics of the motor system and the task at hand during exploration and learning (Sternad, 2018). While these factors certainly contribute to movement variability, our experiments show that a significant portion of the arm’s position, force, and acceleration variability can be predicted by variable timing in the muscles executing the movement.

While chunking and more efficient neural representations (Krakauer et al., 2019; Rhodes et al., 2004) can explain why movements can be executed faster, they do not explain how movements become less variable. Reducing the TV of the muscles involved in a movement appears to be the key to skill learning as less variable movements can be made at the same speed (Diedrichsen and Kornysheva, 2015) (**Figure 1F, 2C, and 3D**, see Figure S2 in Supplementary Materials for improvements in speed-precision trade-off during simulated reaching as TV decreases). This difference in movement variability for the same speed has been reported between the dominant and non-dominant hands (Todor and Doane, 1978; Woodworth, 1899). The second experiment showed how the arm whose muscles had a smaller TV produced a periodic force with less variability, and this happened to be the dominant right-hand. We hypothesized that there may be a relationship between handedness and the relative TV of the left and right arms. In the third experiment, we developed an acceleration variability index that estimated the TV of the arm’s muscles via the arm’s acceleration variability during circle tracing. The acceleration variability was smaller in the dominant hand of both left-handers and right-handers, and the difference in the left- and right-hand’s acceleration variability was related to the Edinburgh Quotient, suggesting that the difference in the left and right arms’ TV could be related to handedness. While there may be other factors that determine the difference in the control of the left and right hands, the dominant muscles’ smaller TV could be a significant factor that determines its superior precision.

A number of studies have proposed precise timing to be a key ingredient to coordinating movements (Calvin, 1983; Manto et al., 2012), yet the effects of timing volatility on movement kinematics and dynamics have not been scrutinized. Recent work suggests that timing may be encoded in the population dynamics of neurons or in their state trajectories (Karmarkar and Buonomano, 2007). These networks, which can be trained to produce movements (Laje and Buonomano, 2013), could be partially distributed in cortical areas such as the premotor, motor and supplementary motor areas because transcranial magnetic stimulation aimed at these areas disrupts the execution of muscle activity without affecting its shape or amplitude (Churchland et al., 2006; Day et al., 1989). Additionally, some of these cortical areas also receive inputs from the basal ganglia, thalamus, and cerebellum. The cerebellum is important in sensing and producing timed responses (Manto et al., 2012), and cerebellar lesions cause muscles to activate too early or late during voluntary movements (Perrett et al., 1993). Therefore, the population dynamics needed to encode movement timing may be distributed over the cerebrum, diencephalon, brain stem and the cerebellum. While it is still difficult at this stage to decompose the neural sources that determine the volatility of the commands reaching the muscles, the CTV theory clarifies the computational mechanism of the central origin of movement variability (Churchland et al., 2006; Osu et al., 2015).

If the control of movement timing is widespread across the central nervous system, neurological damage to the above-mentioned areas could impair motor timing. The simulations we conducted suggest that the motor coordination of reaching movements becomes impaired in a certain way if the muscles controlling the arm have extremely volatile timing. Interestingly, while Huntington’s disease, Parkinson’s disease and cerebellar disorders have differing pathologies and affect different neural pathways, they are all known to impair motor timing during finger tapping (Michell et al., 2008; O’Boyle et al., 1996; Schlerf et al., 2007), possibly through degeneration (Lemoine et al., 2021) or dopamine loss in the striatum (Kish et al., 1988), which receives rich inputs from the cerebellum (Kameda et al., 2023). This deficit in the control of motor timing could be a sign of large TV. When we simulated reaching movements with large TV, the trajectory became jerky towards the end of the movement, and large movement variability was observed soon after movement onset. These simulation results are consistent with observations of reaching movements in patients with HD (Smith et al., 2000), PD (Sheridan et al., 1987) and CA (Sanguineti et al., 2003). Other studies have proposed that these deficits during reaching could be due to reduced sensory processing (Boecker et al., 1999) or faulty feedback error control (Smith et al., 2000), but neither of these theories can explain why the movement variability of the patients is so large at the beginning of the movement where mostly feedforward control is involved (Kawato, 1999). The CTV theory, on the other hand, reveals how increasing TV magnifies movement variability in the beginning and the end of the movement in the form of impaired aiming and jerkiness, thereby impairing both feedforward and feedback control.

The CTV theory proposes a novel computational perspective of how seemingly distinct motor phenomena, from handedness to some motor symptoms of neurodegeneration, may have a common underlying factor that is swayed by the volatility in the timing of the muscles. While the CTV theory fills major gaps in our understanding of motor control, it also raises many new questions. How does the brain learn a neural representation that minimizes timing volatility? Which cortical and sub-cortical loops determine timing volatility? Do reductions in command timing volatility transfer to other tasks, muscles, and limbs? These and many more avenues of research are opened by the insight that command timing volatility plays a significant role in determining the variability of movements.

## Methods

### Simulation of force variability during single-joint reaching movement

To simulate the force variability during single-joint reaching movements caused by different sources of noise in the periphery and in the movement planning stage as predicted by the SDN, SN, TN and CTV theories (**Figure 1A**), we emulated a movement composed of a burst in agonist followed by antagonist muscle activity. The muscle activity at time *t* had a Gaussian shape with the form

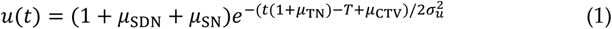

where σ*_u_* = 70 ms and the timing of the peak muscle activity *T* was set to *T* = 200 ms for the flexor and *T* = 400 ms for the extensor muscles. The variables *μ* were noise variables sampled from a Gaussian distribution with σ_SDN_ = 0.25, σ_SN_ = 0.25, σ_TN_ = 0.1 and σ_CTV_ = 50 ms. The noise was sampled once per simulated movement. *μ*_SN_ and *μ*_TN_ were the same for the agonist and antagonist muscles, but *μ*_SDN_ and *μ*_CTV_ were sampled separately for each muscle. To predict the force variability from one theory, the noise variables from other theories were set to zero. For example, to simulate the effect of SDN on the force variability, *μ*_SDN_ was varied but *μ*_SN_ = 0, *μ*_TN_ = 0 and *μ*_CTV_ = 0.

In this simulation, we were more concerned about the shape of the force variability time-series rather than its amplitude, so the standard deviation of the noise values were chosen to have comparable force variability time-series amplitudes for comparison purposes in **Figure 1A**.

### Experiments

All three experiments were performed at NTT Communication Science Laboratories, Japan. The studies were approved by the institutional ethics committee with all participants providing written informed consent prior to participation (no. R05-014 for first two experiments and R03-011 for the third).

### Elbow reaching experiment

Ten right-handed participants (all male, aged 35±3) made 50 elbow reaching movements using their left arm by starting from [0.01, 0.18] m and reaching towards a target placed at [0.11, 0.04] m using a robotic manipulandum (KINARM endpoint, BKIN Technologies). Participants rested their elbow on an aluminium beam to ensure only the elbow was used in the task. Feedback of movement duration was given to participants after every trial if the duration was outside the range between 400-500 ms (‘slow’ or ‘fast’). The timer started when the participant left the starting circle of radius 1 cm and ended when their velocity was lower than 0.1 m/s and their position was inside the target circle of radius 1.5 cm. Surface electromyograms (EMG) of the biceps brachii and the triceps lateral head were recorded using wireless electrodes (picoEMG, Cometa). The skin was cleaned using alcohol pads prior to the application of the electrodes. Kinematic and EMG data were all recorded at 1000 Hz. EMG was high-pass filtered using a second-order Butterworth filter with a cutoff of 10 Hz, rectified, then low-pass filtered at 9 Hz to obtain the envelope. All 50 movements were used in the analysis irrespective of whether the movement duration was within or outside the prescribed range.

### Isometric force control experiment

Ten right-handed participants (all male, aged 32±4) with no known neuromuscular disorders or recent injuries took part in the experiments. The Edinburgh Inventory was used to confirm the right-handedness of our participants (Oldfield, 1971). Participants exerted a periodic force against the KINARM robotic interface using either their wrist, elbow, or shoulder. The other joints were fixed with a brace when not being tested. The position of the handle was held strongly at the origin of the task [*x*_0_, *y*_0_]*^T^* using a position controller with a stiffness of 1500 N/m. Participants pushed against this handle to produce a force at a frequency determined by a metronome. The applied force was displayed as a white arrow on a computer monitor facing downwards, which was viewed through a film mirror placed 10 cm above the robotic planar workspace. The mirror blocked direct view of the robot and the entire arm. A 10 N force corresponded to an arrow length of 5 cm onscreen. Two circular targets were always displayed during a trial, which were 6 N away from the centre.

A piezoelectric speaker was connected to the analog output of the robotic interface to produce beeps at a constant frequency. The metronome was placed directly in front of the participant’s head at their midline. Participants were asked to push against the handle periodically to hit each target with the arrow after each beep, hence the length of two beeps corresponded to one period of the arm’s periodic force output. Each frequency was tested consecutively for three trials, and each frequency was tested in the order of {3, 4, 5} Hz. Each trial lasted 6 seconds. The trials were ordered such that the right wrist was tested at all frequency levels first (9 trials), followed by the right elbow, and then the shoulder. Afterwards, participants completed the same procedure using the left wrist, left elbow and left shoulder. Each participant completed 54 trials in total, half using the left and the other half with the right arm.

EMG from the wrist, elbow, and shoulder flexors (flexor carpi radialis, brachioradialis and pectoralis major) and extensors (extensor carpi radialis longus, lateral head of triceps brachii and posterior deltoid) from the left and right arms were measured using wireless electrodes. The EMG data was filtered using the exact same method and parameters as in the elbow reaching experiment. The data of the robotic handle’s position, the force exerted by the participants, and the muscle activity from the left and right shoulder were digitized and recorded at 1000 Hz.

### Periodic movement variability experiment

141 participants (90 males, mean age 35.8±0.9) were recruited to partake in the experiment. Participants were instructed to hold a device (Oppo A35) in either their left or right hand with the screen facing up, and rotated it outwards in a circular fashion (clockwise for right hand, anti-clockwise for the left) with a radius of approximately 10 cm. The device produced a periodic beep at 2.5 Hz. Participants had to complete one revolution of the movement between each beep. 15 seconds of data was collected per hand, with the first 1.5 seconds being discarded from the analysis. The data from each hand was analyzed separately. The offset from the acceleration data was removed by subtracting the mean value from each axis. Then the acceleration values were divided by their Euclidean vector norm averaged over the entire 15 second duration to even out the amplitude for between-hand and between-subject comparisons. The data was then segmented into cycles, where each cycle started and ended when the angle of the acceleration along the x and y axes crossed 0°. The Hausdorff distance between two neighboring cycles was calculated beginning from the first cycle until the end, and the mean distance from all neighboring cycle pairs was defined to be σ*_a_*.

### Simulation of the brain controlling a two-joint arm

The arm was modelled as a two-joint, six muscle system with the shoulder’s Cartesian position fixed in place. The shoulder and elbow joints each had two monoarticular muscles that pulled in opposite directions, and two biarticular muscles exerted a torque simultaneously on the shoulder and elbow joints. Each muscle was modelled as a linear spring-damper that connected the joint segment to its corresponding joint limit angle such that its viscoelasticity increased linearly with muscle activation, generating a restoring force to the neutral posture which depended on the activity of all muscles spanning that joint (Takagi et al., 2022) (see Supplementary Materials for details). The six muscles altogether exerted a torque ***τ*** on the arm such that its state **q** = [*q_s_*, *q_e_*]^′^ evolved in time according to

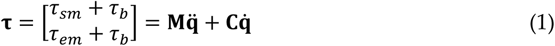

where *τ_sm_*, *τ_em_* and *τ_b_* were the torques from the shoulder monoarticular, elbow monoarticular and biarticular muscles (see Supplementary Materials for details) and the arm’s inertia in joint space

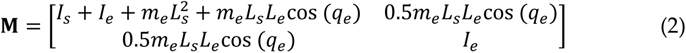

was a function of the elbow angle and

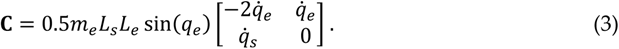

was the Coriolis force. The parameters *L_s_* = 0.3 m and *L_e_* = 0.35 m are the lengths of the shoulder and elbow segments, *I_s_* = 0.05 kgm^2^ and *I_e_* = 0.06 kgm^2^ are their moments of inertia, and *m_s_* = 1.6 kg and *m_e_* = 1.4 kg are their point masses (Katayama and Kawato, 1993). The joint inertia assumes that the location of the elbow’s center of mass is halfway along its segment.

This arm was controlled using Model Predictive Control, which has been used to accurately predict multi-joint human movements (Da Silva et al., 2008; Takagi et al., 2022; Teramae et al., 2018), by planning the muscle activity needed to reach the target within some receding horizon *T* = 150 ms. If the target is not reached after *T* seconds, MPC replanned the movement until the target was reached. The arm’s state **ξ** = [**q**, q̇]^′^ was composed of the two-link arm’s joint positions and velocities, and the goal was to move this arm to a new position and a set velocity **ξ_*_** using the muscle activity **u**. Defining time to be *t*, then the cost function *W* took the form

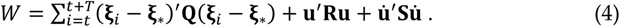

The first term, which is weighted by **Q** = Diag(300, 300, 4, 4), minimized the distance between joint’s predicted state **ξ***_i_* and the desired state **ξ_*_**. The second and third terms, weighted by **R** and **S** respectively, punished unnecessary muscle activity and rapid changes in muscle activity to conserve energy (Wong et al., 2021). These weights were the same for all muscles, rendering them as scalar constants *R* = 10^−5^ and *S* = 3. While the muscle activity was constant for the horizon, any change in the joint’s position or its velocity resulted in a change in joint torque in accordance with eq. (1). **Q** = Diag(300, 300, 4, 4), *R* = 10^−5^, *S* = 3.

MPC predicted the joint state **ξ** iteratively by starting from its last measured value and iterating it forwards in time using the dynamic model **ξ***_i_*_+1_ = **f**(**ξ***_i_*, **u**). We used a parametric model for the internal model **f**, and the Euler method was used to update the state,

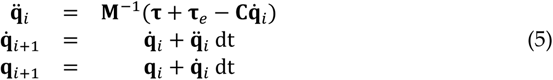

The optimization problem was to find the muscle activity **u ≥ 0** that minimized the cost function in eq. (4) using eq. (1). This was solved by Quadratic Programming (QP) or by iteratively updating the muscle activity until the solution converged to an acceptable minimum. We used the interior-point method implemented in MATLAB’s ‘fmincon’ function (using its default settings) to solve this problem due to its speed and robustness.

Every instance that *T* seconds elapsed, a new motion plan with a new muscle activity **u** was found. The planned muscle activity was filtered by a first-order low pass filter with a time constant of 50 milliseconds to reproduce the filter-like properties of the muscle (Winter, 2009). All simulations were discretized with a time step of *dt* = 0.01 s. The arm’s initial posture was specified for each interaction scenario simulation. Non-zero muscle activity was needed to sustain the arm at a given initial posture, and so ‘fmincon’ was used to find the initial muscle activity that produced zero torque at a given initial joint posture.

Timing volatility on the arm’s control was implemented by sending the planned muscle activity **u** either ahead or late in time to each muscle. If the ideal timing to send the muscle activity was *t* = 0, then the actual time was sampled from white noise with standard deviation σ*_t_* = 80, 160, 240 or 320 ms. The timing noise from a pair of flexor and extensor muscles spanning the same joint were sampled to have a correlation of 0.15, the approximate value measured from the second experiment.

### Statistical analysis

T-tests were used to examine the significance of the difference in the force variability and TV in the left and right hands in the isometric force control experiment. Holm-Bonferroni corrections were applied to control for multiple comparisons.

Robust regression was used to assess the significance of the slope between the elbow reaching variability and TV, and it was used was used to check the significance of the slope between elbow reaching variability and SDN or variations in the agonist and antagonist’s size variability.

Linear mixed-effects analysis was used to assess the significance of the linear relationship between TV and the force variability in the isometric experiment, and between TV estimated using the smartphone and TV measured by the EMG during the isometric experiment.

A two-way repeated measures ANOVA with the hand (left, right) and group (left-hander, right-hander, forced-hander) as the main factors was used to test for differences in the TV between the left and right hands amongst the three groups, and post-hoc tests were controlled for multiple comparisons using Tukey’s HSD.

## Acknowledgements

The authors would like to thank Mitsuo Kawato for commenting on an earlier draft. A.T. was partially supported by the JST PRESTO grant JPMJPR18J5. H.G. was partially supported by Grants-in-Aid for Scientific Research (JP16H06566) from Japan Society for the Promotion of Science.

## References

Adam A, Luca CJD, Erim Z. 1998. Hand Dominance and Motor Unit Firing Behavior. J Neurophysiol 80:1373–1382. doi:10.1152/jn.1998.80.3.1373

Beers RJ van, Sittig AC, Gon JJD van der. 1999. Integration of Proprioceptive and Visual Position-Information: An Experimentally Supported Model. J Neurophysiol 81:1355–1364.

Berret B, Jean F. 2020. Stochastic optimal open-loop control as a theory of force and impedance planning via muscle co-contraction. PLOS Comput Biol 16:e1007414. doi:10.1371/journal.pcbi.1007414

Boecker H, Ceballos-Baumann A, Bartenstein P, Weindl A, Siebner HR, Fassbender T, Munz F, Schwaiger M, Conrad B. 1999. Sensory processing in Parkinson’s and Huntington’s disease: Investigations with 3D H215O-PET. Brain 122:1651–1665. doi:10.1093/brain/122.9.1651

Calvin WH. 1983. A stone’s throw and its launch window: Timing precision and its implications for language and hominid brains. J Theor Biol 104:121–135. doi:10.1016/0022-5193(83)90405-8

Churchland MM, Afshar A, Shenoy KV. 2006. A Central Source of Movement Variability. Neuron 52:1085–1096. doi:10.1016/j.neuron.2006.10.034

Da Silva M, Abe Y, Popović J. 2008. Simulation of Human Motion Data using Short-Horizon Model-Predictive Control. Comput Graph Forum 27:371–380. doi:10.1111/j.1467-8659.2008.01134.x

Day BL, Rothwell JC, Thompson PD, Maertens de Noordhout A, Nakashima K, Shannon K, Marsden CD. 1989. Delay in the execution of voluntary movement by electrical or magnetic brain stimulation in intact man. Evidence for the storage of motor programs in the brain. Brain J Neurol 112 ( **Pt 3**):649–663. doi:10.1093/brain/112.3.649

Diedrichsen J, Kornysheva K. 2015. Motor skill learning between selection and execution. Trends Cogn Sci 19:227–233. doi:10.1016/j.tics.2015.02.003

Gordon J, Ghilardi MF, Ghez C. 1994. Accuracy of planar reaching movements. Exp Brain Res 99:97–111. doi:10.1007/BF00241415

Hansen RM, Skavenski AA. 1977. Accuracy of eye position information for motor control. Vision Res 17:919–926. doi:10.1016/0042-6989(77)90067-0

Harris CM, Wolpert DM. 1998. Signal-dependent noise determines motor planning. Nature 394:780–784. doi:10.1038/29528

Huttenlocher DP, Klanderman GA, Rucklidge WJ. 1993. Comparing images using the Hausdorff distance. IEEE Trans Pattern Anal Mach Intell 15:850–863. doi:10.1109/34.232073

Jones KE, Hamilton AF de C, Wolpert DM. 2002. Sources of Signal-Dependent Noise During Isometric Force Production. J Neurophysiol 88:1533–1544. doi:10.1152/jn.2002.88.3.1533

Kameda M, Niikawa K, Uematsu A, Tanaka M. 2023. Sensory and motor representations of internalized rhythms in the cerebellum and basal ganglia. Proc Natl Acad Sci 120:e2221641120. doi:10.1073/pnas.2221641120

Karmarkar UR, Buonomano DV. 2007. Timing in the Absence of Clocks: Encoding Time in Neural Network States. Neuron 53:427–438. doi:10.1016/j.neuron.2007.01.006

Katayama M, Kawato M. 1993. Virtual trajectory and stiffness ellipse during multijoint arm movement predicted by neural inverse models. Biol Cybern 69:353–362. doi:10.1007/BF01185407

Kawato M. 1999. Internal models for motor control and trajectory planning. Curr Opin Neurobiol 9:718–727. doi:10.1016/S0959-4388(99)00028-8

Kish SJ, Shannak K, Hornykiewicz O. 1988. Uneven Pattern of Dopamine Loss in the Striatum of Patients with Idiopathic Parkinson’s Disease. N Engl J Med 318:876–880. doi:10.1056/NEJM198804073181402

Krakauer JW, Hadjiosif AM, Xu J, Wong AL, Haith AM. 2019. Motor learning. Compr Physiol 9:613–663. doi:10.1002/cphy.c170043

Laje R, Buonomano DV. 2013. Robust timing and motor patterns by taming chaos in recurrent neural networks. Nat Neurosci 16:925–933. doi:10.1038/nn.3405

Lemoine L, Lunven M, Bapst B, Cleret de Langavant L, de Gardelle V, Bachoud-Lévi A-C. 2021. The specific role of the striatum in interval timing: The Huntington’s disease model. NeuroImage Clin 32:102865. doi:10.1016/j.nicl.2021.102865

Manto M, Bower JM, Conforto AB, Delgado-García JM, da Guarda SNF, Gerwig M, Habas C, Hagura N, Ivry RB, Mariën P, Molinari M, Naito E, Nowak DA, Oulad Ben Taib N, Pelisson D, Tesche CD, Tilikete C, Timmann D. 2012. Consensus Paper: Roles of the Cerebellum in Motor Control—The Diversity of Ideas on Cerebellar Involvement in Movement. The Cerebellum 11:457–487. doi:10.1007/s12311-011-0331-9

Meyer DE, Smith JE, Wright CE. 1982. Models for the speed and accuracy of aimed movements. Psychol Rev 89:449–482.

Michell AW, Goodman AOG, Silva AHD, Lazic SE, Morton AJ, Barker RA. 2008. Hand tapping: A simple, reproducible, objective marker of motor dysfunction in Huntington’s disease. J Neurol 255:1145–1152. doi:10.1007/s00415-008-0859-x

O’Boyle DJ, Freeman JS, Cody FWJ. 1996. The accuracy and precision of timing of self-paced, repetitive movements in subjects with Parkinson’s disease. Brain 119:51–70. doi:10.1093/brain/119.1.51

Oldfield RC. 1971. The assessment and analysis of handedness: The Edinburgh inventory. Neuropsychologia 9:97–113. doi:10.1016/0028-3932(71)90067-4

Osu R, Morishige K, Nakanishi J, Miyamoto H, Kawato M. 2015. Practice reduces task relevant variance modulation and forms nominal trajectory. Sci Rep 5:17659. doi:10.1038/srep17659

Perrett SP, Ruiz BP, Mauk MD. 1993. Cerebellar cortex lesions disrupt learning-dependent timing of conditioned eyelid responses. J Neurosci 13:1708–1718. doi:10.1523/JNEUROSCI.13-04-01708.1993

Rhodes BJ, Bullock D, Verwey WB, Averbeck BB, Page MPA. 2004. Learning and production of movement sequences: Behavioral, neurophysiological, and modeling perspectives. Hum Mov Sci, European Workshop on Movement Science 23:699–746. doi:10.1016/j.humov.2004.10.008

Sanguineti V, Morasso PG, Baratto L, Brichetto G, Luigi Mancardi G, Solaro C. 2003. Cerebellar ataxia: Quantitative assessment and cybernetic interpretation. Hum Mov Sci, Advances in the Study of Drawing and Handwriting 22:189–205. doi:10.1016/S0167-9457(02)00159-8

Schlerf JE, Spencer RMC, Zelaznik HN, Ivry RB. 2007. Timing of rhythmic movements in patients with cerebellar degeneration. The Cerebellum 6:221–231. doi:10.1080/14734220701370643

Sheridan MR, Flowers KA, Hurrell J. 1987. Programming and execution of movement in Parkinson’s disease. Brain J Neurol 110 (Pt 5):1247–1271. doi:10.1093/brain/110.5.1247

Shmuelof L, Krakauer JW, Mazzoni P. 2012. How is a motor skill learned? Change and invariance at the levels of task success and trajectory control. J Neurophysiol 108:578–594. doi:10.1152/jn.00856.2011

Smith MA, Brandt J, Shadmehr R. 2000. Motor disorder in Huntington’s disease begins as a dysfunction in error feedback control. Nature 403:544–549. doi:10.1038/35000576

Sternad D. 2018. It’s not (only) the mean that matters: variability, noise and exploration in skill learning. *Curr Opin Behav Sci*, Habits and Skills 20:183–195. doi:10.1016/j.cobeha.2018.01.004

Takagi A, Gomi H, Burdet E, Koike Y. 2022. A model predictive control strategy to regulate movements and interactions. doi:10.1101/2022.08.24.505193

Teramae T, Noda T, Morimoto J. 2018. EMG-Based Model Predictive Control for Physical Human–Robot Interaction: Application for Assist-As-Needed Control. IEEE Robot Autom Lett 3:210–217. doi:10.1109/LRA.2017.2737478

Todor JI, Doane T. 1978. Handedness and Hemispheric Asymmetry in the Control Of Movements. J Mot Behav 10:295–300. doi:10.1080/00222895.1978.10735163

van Beers RJ, Haggard P, Wolpert DM. 2004. The Role of Execution Noise in Movement Variability. J Neurophysiol 91:1050–1063. doi:10.1152/jn.00652.2003

Winter DA. 2009. Biomechanics and Motor Control of Human Movement. John Wiley & Sons.

Wong JD, Cluff T, Kuo AD. 2021. The energetic basis for smooth human arm movements. eLife 10:e68013. doi:10.7554/eLife.68013

Woodworth RS. 1899. Accuracy of voluntary movement. Psychol Rev Monogr Suppl 3:i–114. doi:10.1037/h0092992

Zimnik AJ, Churchland MM. 2021. Independent generation of sequence elements by motor cortex. Nat Neurosci 24:412–424. doi:10.1038/s41593-021-00798-5

