## Supplementary Materials for "Minimizing command timing volatility is a key factor in skilled actions"

Simulations were conducted on a single-joint model of the arm (elbow only) to examine the effect of muscular cocontraction on the position variability caused by TV, and to examine the relationship between peak movement speed and movement variability from TV. The model parameters were identical to those used in the Methods.

The initial elbow angle was set to $q_{e}=83$ deg, and the target angle was set to 130 deg to produce elbow movements of amplitude 15 cm. The cost matrices were set to $\mathbf{Q}=\mathrm{diag}(2000, 70)$, $R=3$and $S=10$. Movement speed was varied by changing the velocity-term of the cost matrix **Q** between [10, 120] in steps of 11. The cocontraction level was set by constraining the Model Predictive Controller (MPC) to produce a minimum muscle activity **u** equal to the cocontraction level $u_{c}$. This forced MPC to solve the optimization problem and generate movements that used a minimum cocontraction $u_{c}$. 200 reaching movements were simulated per speed or cocontraction level.

When reaching with different amounts of cocontraction, the position and force time-series did not change significantly (Figure S2A, compare solid and dashed traces). However, the position and force variabilities were dependent on the level of cocontraction, and this dependency was different for SDN and CTV. With SDN, cocontraction led to an increase in the position and force variability, while the opposite effect was observed with CTV. We calculated the mean position variability at the end of the movement as a function of the change in overall muscle activity, which was normalized to be 100% when $u_{c}=0$ (Figure S2B). A gradual increase in cocontraction caused the position variability due to SDN to increase alongside it. On the other hand, the movement variability due to TV decreased with increasing cocontraction, which is the expected behavior.

A major selling point of the SDN theory is its ability to reproduce Fitts’ law or the linear relationship between speed and accuracy. We calculated the position variability due to TV under different movement speeds (Figure S2C). The CTV theory also reproduces Fitts’ law, and different amounts of TV resulted in a change in the speed-accuracy trade-off, implying that reducing TV is the key to reaching with high precision at the same speed, i.e., skill learning.

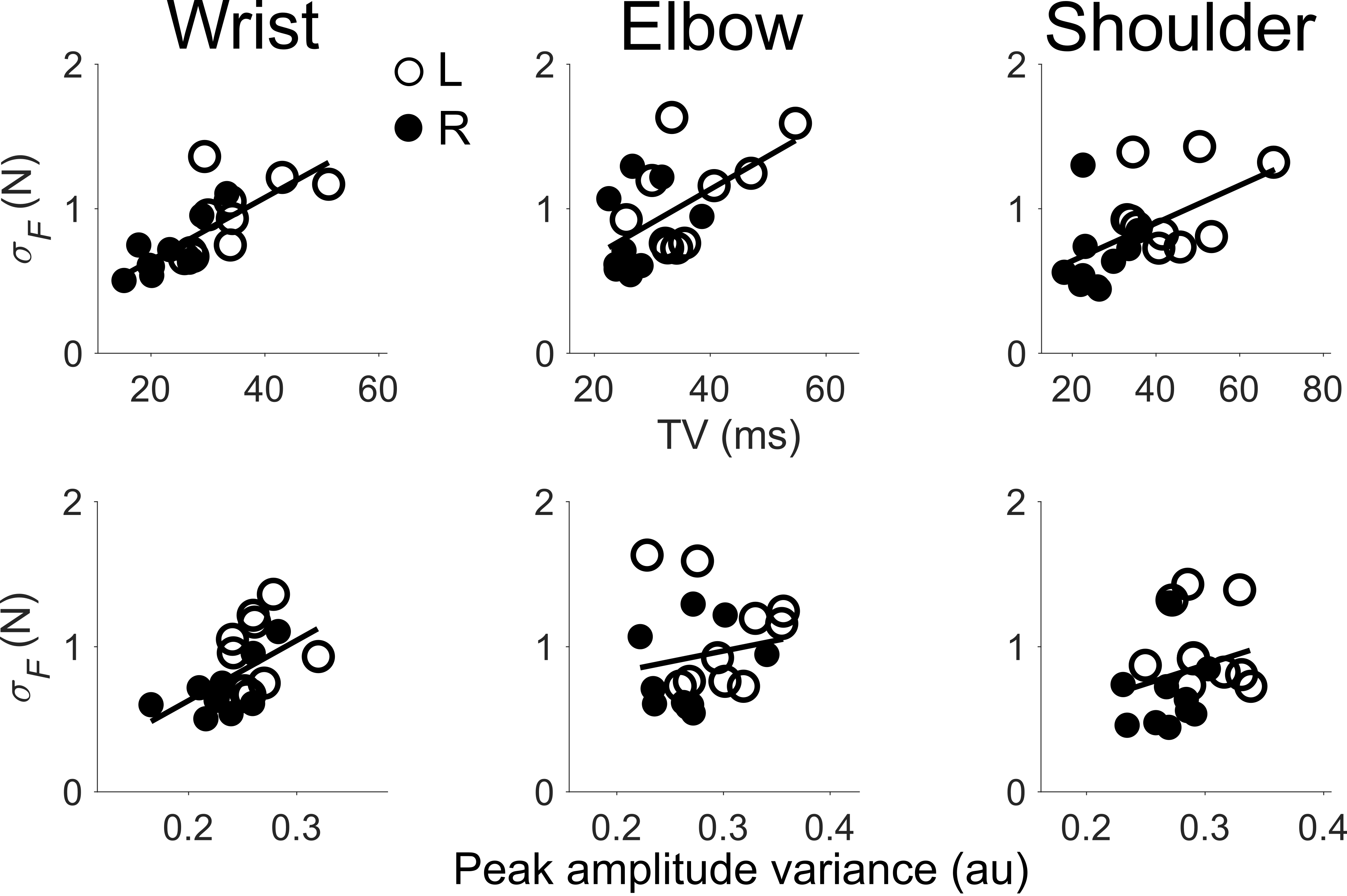

**Figure S1.** Data from the second isometric force control experiment where participants exerted a periodic force isometrically using either their wrist, elbow, or shoulder joints from their left (open circle) or right arm (filled). A linear mixed-effects analysis was conducted with the force variability as the dependent variable and the TV, the variations in the peak EMG size, the categorial hand (left, right) and categorical joint (wrist, elbow, shoulder) variables as the predictors. This revealed no effect of the joint factor, so it was collapsed in Figure 2C, but here it has been separated to show how the relationship between force variability and TV can be observed in all three joints. (Top row) Timing error in the agonist and antagonist muscles (TV) as a function of the force variability. (Bottom row) Variations in the size of the peak EMG as a function of force variability.

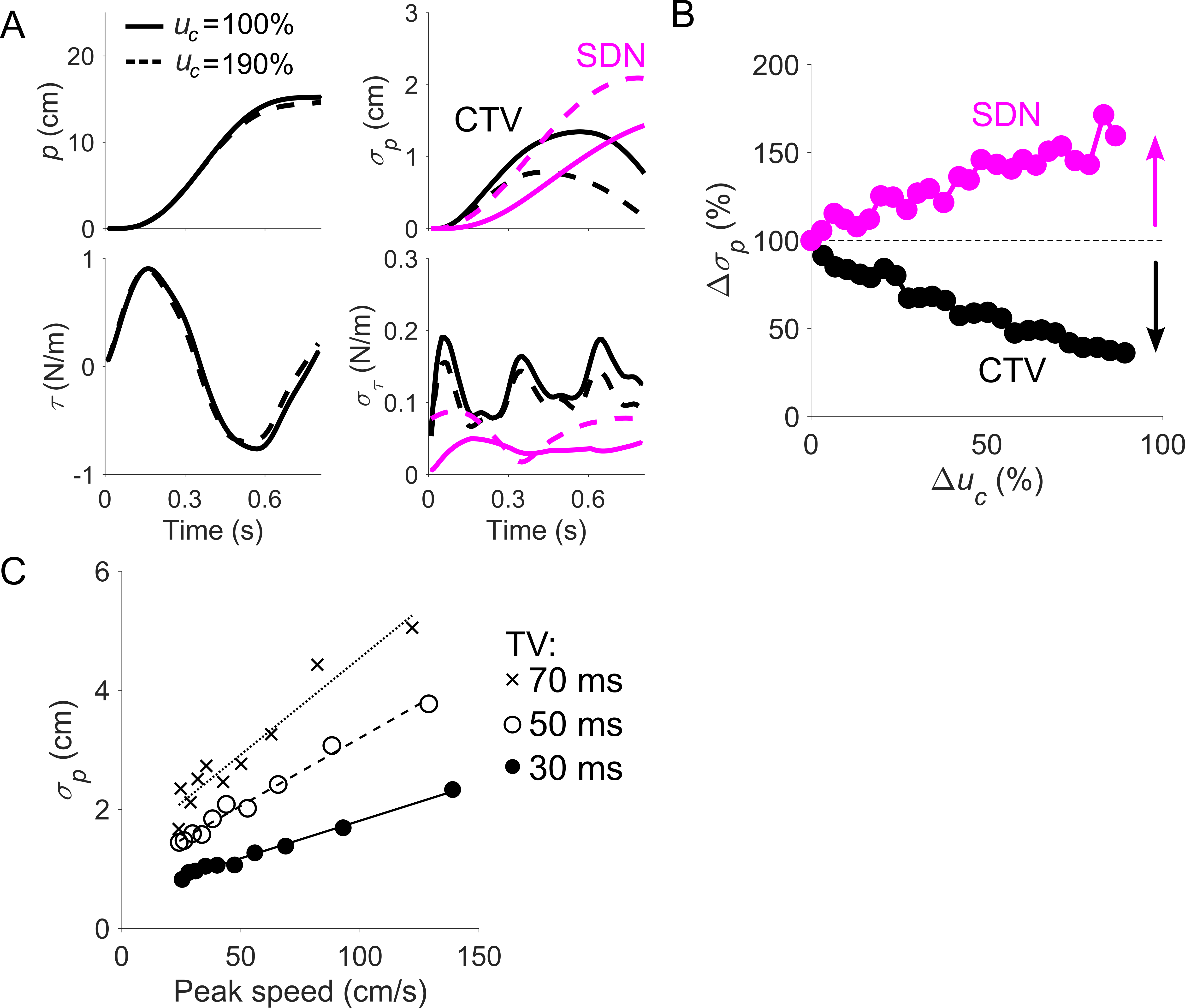

**Figure S2.** Position variability produced by TV decreases due to cocontraction, and the relationship between movement speed and position variability due to TV follows Fitt’s law. (A) Position and force time-series did not change significantly when reaching with different levels of cocontraction, but their variability did. Movement variability from SDN increased with larger cocontraction while the opposite trend was observed for timing volatility. (B) Larger cocontraction increased position variability due to SDN, whereas the opposite occurred for TV. (C) TV reproduces Fitt’s law or the linear relationship between speed and precision. A change in TV shifts the speed-accuracy trade-off, implying that skill learning is achieved by minimizing TV.
